# Mutant SOD1 expressed by oligodendrocytes aggregates in myelinic nanochannels and accelerates disease progression in familial ALS mice

**DOI:** 10.64898/2026.06.09.731100

**Authors:** Alexandra I. Mot, Ying Li, Payam Dibaj, Iva D. Tzvetanova, Ulrike C. Gerwig, Tizibt A. Bogale, Sandra Goebbels, Wiebke Möbius, Dwight E. Bergles, Brett M. Morrison, Jeffrey D. Rothstein, Don W. Cleveland, Julia M. Edgar, Klaus-Armin Nave

**Author notes:** Corresponding authors: Don W. Cleveland, Klaus-Armin Nave.

## Abstract

Amyotrophic lateral sclerosis (ALS) is a highly debilitating and fatal disease characterized by the progressive loss of motor neurons. Reduced oligodendroglial support has been implicated in ALS progression but remains mechanistically unexplained. Here, using a mutant superoxide dismutase 1 (SOD1-G37R) mouse model of familial ALS, Cre-mediated excision of the mutant SOD1 gene within the oligodendrocyte lineage prior to myelin compaction is shown to slow disease onset, improve motor performance, and prolong survival. In contrast, silencing mutant *SOD1* expression within oligodendrocytes after myelin compaction failed to ameliorate disease phenotype. Electron microscopy is used to identify aggregation of mutant SOD1 within paranodal loops and the inner periaxonal tongue of ‘myelinic nanochannels’, narrow cytosolic compartments for the diffusion of metabolites and motor-driven transport processes. In a second mouse model (SOD1-G93A) of familial, SOD1 mutant-mediated ALS, we show that induction of excessive myelin compaction and myelinic channel collapse (by depletion of CNP from myelin) accelerates disease and diminishes survival. Our data support loss of myelinic channel integrity as a contributor to familial ALS disease initiation and progression, findings likely relevant to neurodegenerative disease involving other aggregation prone proteins that are expressed in myelinating oligodendrocytes.

**Significance Statement:** Oligodendrocytes have been implicated in the progression of amyotrophic lateral sclerosis (ALS) but the underlying mechanisms have remained obscure. Here we show in genetic mouse models that the familial ALS causing isoform of a ubiquitously expressed mutant enzyme (SOD1) aggregates in cytosolic channels within myelin that are responsible for delivery of transporters and nutrients necessary to support the axonal compartment. ALS disease progression was accelerated in mice when myelinic channels were collapsed by deleting CNP, a structural protein necessary for myelinic channel maintenance. Disruption of transport through myelinic channels by aggregation of mutant SOD1 may perturb oligodendrocyte support of motor axons and contribute to disease in this form of ALS.

## Introduction

Amyotrophic lateral sclerosis (ALS) is the most common form of adult motor neuron disease. Although the majority of ALS cases are sporadic, about 10% are caused by genetic mutations (*1*). Most widely studied are those associated with ubiquitously expressed ALS genes, such as superoxide dismutase 1 (*SOD1*), transactive response DNA binding protein 43 (*TDP43*), fused in sarcoma (*FUS*), and hexanucleotide repeat expansions in chromosome 9 open reading frame 72 (*C9ORF72*). *SOD1*, the first gene identified in familial ALS has become the prototype also due to the availability of transgenic models that exhibit reproducible pathology (*2*). The encoded enzyme plays a central role in combating oxidative stress in cells and the mutant protein is thought to contribute to neuronal death by a toxic gain-of-function mechanism (*3*). Although ALS is primarily characterized by the loss of motor neurons, disease progression is driven by mutant SOD1 synthesis within astrocytes (*4*) and microglia (*5*), thereby producing non-cell autonomous pathogenesis by the ubiquitously expressed mutant SOD1 (*6, 7*).

A specific feature of motor neurons are their long axonal projections with high energy demands. Myelinating oligodendrocytes provide motor neurons with metabolic support (*8*). The fact that motor axons are almost completely encapsulated by the myelin sheath significantly may limit their access to extracellular metabolites, making direct metabolic support by oligodendrocytes more important (*9*). Transport of oligodendroglial glycolysis products through the myelin sheath to the glial-axonal junction occurs via a network of narrow cytosolic channels, spanning 20 to 300 nanometer in width and including outer/inner tongues of myelin and the paranodal loops, collectively referred to as “myelinic nanochannels” (*10, 11*). In addition to direct glycolytic support (*12, 13*), oligodendrocytes also release exosomes, in an activity-dependent manner, whose contents are taken up by axons beneath the myelin sheath (*14, 15*). Here, the transfer of ferritin heavy chain to axons provides antioxidant defense (*16*) whereas SIRT2, an abundant NAD-dependent deacetylase in oligodendrocytes (*17*), activates the mitochondrial generation of ATP in the myelinated axon (*18*). Most neurodegenerative diseases, including ALS exhibit increasing pathology with age; however, less is known about how progressive changes in oligodendrocytes may contribute to neuronal dysfunction (*19*).

Recent studies reported that oligodendroglial dysfunction plays a role in ALS pathogenesis, as reviewed in (*20*). Progressive demyelination of the spinal cord in both sporadic and familial ALS patients, as well as in SOD1-G93A mutant mice (*21*) has been observed. In response to damage to and turnover of oligodendrocytes, there is increased proliferation of oligodendrocyte progenitor cells (OPCs), but newly formed oligodendrocytes fail to mature, ultimately resulting in sustained demyelination (*22, 23*). In the SOD1-G93A mutant mouse spinal cord newly generated oligodendrocytes are often associated with degenerating axons (*21*), a finding compatible with an oligodendroglial failure of axon support. Moreover, human oligodendrocytes, derived from induced pluripotent stem cells of ALS patients, have decreased ability to support co-cultured motor neurons, leading to decreased neuronal survival (*24*). Additionally, oligodendrocytes have been reported to provide trophic support and neuroprotection to motor neurons exposed to cerebrospinal fluid from sporadic ALS patients (*25*).

Several studies have identified ALS-linked protein aggregates in the soma of oligodendrocytes, including SOD1, TDP43 and FUS (*24, 26–30*). Although the pathogenic consequences of these protein accumulations remain obscure, selective removal of mutant *SOD1* expression from OPCs in SOD1-G37R transgenic mice delays disease onset and prolongs survival (*21*), suggesting that these changes ultimately impair motor neuron survival.

Here, we show that aggregation of mutant SOD1 occurs in myelinic nanochannels, with silencing of mutant *SOD1* expression in oligodendrocytes before myelin compaction producing delayed disease onset and progression, improved motor performance, and prolonged survival, while silencing of mutant *SOD1* expression in oligodendrocytes after myelin compaction failed to ameliorate disease phenotype. Moreover, collapse of myelinic nanochannels from silencing expression of CNP, a myelin protein that maintains the structural integrity of these channels, accelerated disease progression.

## Results

### SOD1-G93A aggregates in oligodendrocyte processes and myelinic nanochannels

The ubiquitous expression of SOD1 includes oligodendrocytes, which metabolically support motor axons and are involved in ALS pathology (*13, 21*). To test our hypothesis that mutant SOD1 isoforms physically aggregate in the nanometer-wide cytosolic channels of the myelin sheaths, we visualized SOD1 aggregates in oligodendrocytes derived from SOD1-G93A ALS mice which express high levels of an ALS-causing mutation in human SOD1 encoded by a multicopy transgene under its own promoter (*31*). OPCs were isolated from embryonic brains and differentiated *in vitro* for 12 days to allow mature oligodendrocytes sufficient time for the formation of SOD1 aggregates. Using an antibody specific for misfolded human SOD1 (MediMabs Cat# MM-0070-P, RRID:AB_2909641), the SOD1 protein aggregates were readily identified in oligodendroglial cell bodies (Fig. 1A) derived from SOD1 mutant mice, but never from wildtype controls. Importantly, SOD1 aggregates were also numerous between myelin-like membrane sheets within narrow, non-compacted cell processes. Interestingly, in thicker processes these SOD1 aggregates were also larger (Fig. 1A insert), consistent with their growth to fill the available space and suggesting that SOD1 aggregates that fill the oligodendrocyte processes prior to myelination might later “clog” these myelinic nanochannels.

**Figure 1.**
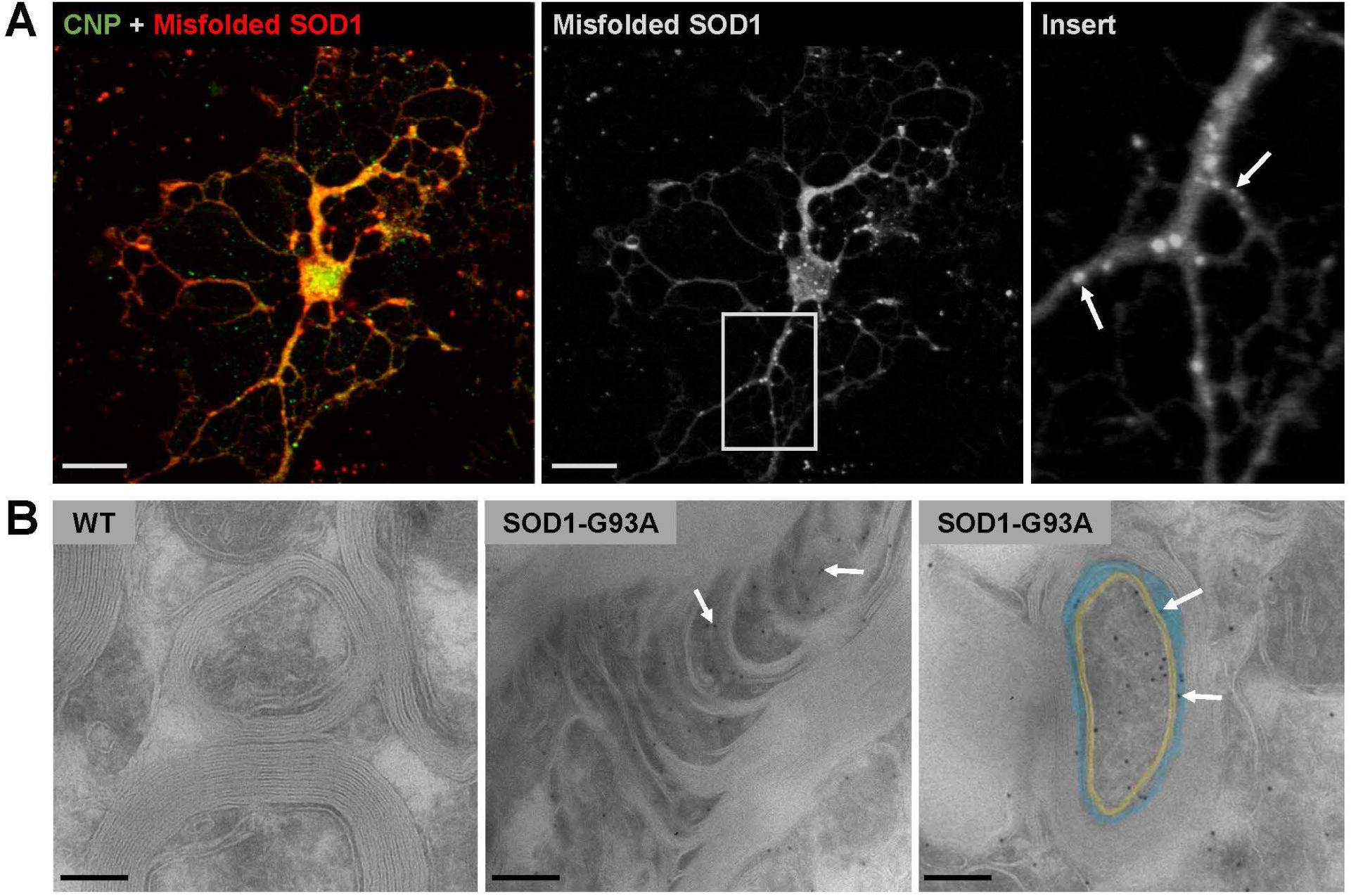
Mutant SOD1 aggregates in oligodendroglial cell processes and myelinic nanochannels. A) Oligodendrocytes derived from SOD1-G93A mutant mouse spinal cords display misfolded SOD1 aggregates in cell processes (insert arrows) *in vitro*. Oligodendrocytes were identified by CNP immunostaining (green) in left panel. Scale bars represent 20µm. B) Imunoelectron micrographs of 5 month old WT and SOD1-G93A spinal cord showing SOD1-gold labelling in non-compacted areas of the myelin sheath including the inner tongue shown in blue (arrows in right panel) and the paranodal loops (arrows in middle panel). The periaxonal space in the right panel is shown in yellow. Scale bars represent 150nm (left and right panels) or 200nm (middle panel).

To identify mutant SOD1 in native myelin, electron microscopy and SOD1 immuno-gold labeling was performed in spinal cord samples collected from 5 months old SOD1-G93A mice. SOD1 was detected in some non-compacted areas of the myelin sheath, including the inner tongue and the paranodal loops, which comprise parts of the myelinic nanochannel network (Fig. 1B). Although the antibody used (Enzo Life Sciences Cat# ADI-SOD-100-D, RRID:AB_2039583) also binds to mouse SOD1, no label was observed in wildtype mice (Fig. 1B), suggesting that wildtype mouse SOD1 is not sufficiently abundant for immunodetection using this method.

### Early removal of mutant SOD1 from the oligodendrocyte lineage

For cell type-specific ablation of ubiquitously expressed mutant SOD, a related ALS mouse model has been generated with a floxed transgene, encoding SOD1-G37R (*4, 5, 21*). We first targeted SOD1-G37R early in the oligodendroglial lineage, i.e. long before myelination begins, using Cre recombinase expression under control of the Sox10 promoter (*32*). *Sox10* is expressed in proliferating OPCs and continues to be expressed in mature oligodendrocytes throughout life (*33*). Crossbreeding with hemizygous loxSOD1^G37R^ mutants yielded four genotypes in the expected Mendelian ratio (Fig. 3A).

To assess the expression of mutant SOD1 in Sox10-Cre; loxSOD1^G37R^ mice, we performed immunohistochemistry and Western blot analyses of the lumbar spinal cord beginning at early disease stages and before motor neuron loss (age 44 weeks). For immunohistochemistry we used an antibody that binds exclusively to misfolded human SOD1 (MediMabs Cat# MM-0070-P, RRID:AB_2909641) and counter-stained spinal cord white matter with an antibody against myelin basic protein (MBP). In lumbar spinal cord sections of Sox10-Cre; loxSOD1^G37R^ mice, there was substantially less SOD1 immunoreactivity when compared to loxSOD1^G37R^ controls (Fig. 2A); there was a mean decrease of 58% in grey matter (Fig. 2B) and a mean decrease of 45% in white matter (Fig. 2C), although in both cases this decrease did not reach statistical significance, due to the high variability between animals (n=4). In both loxSOD1^G37R^ and Sox10-Cre; loxSOD1^G37R^ mice, SOD1 aggregates clustered in the ventral grey matter (insert in Fig. 2A). As expected, there was no misfolded human SOD1 staining in WT or Sox10-Cre controls (Fig. 2A), since these mice only express the endogenous murine *Sod1* gene. For the Western blot analysis an antibody that binds to both human and mouse SOD1 (Enzo Life Sciences Cat# ADI-SOD-100-D, RRID:AB_2039583) was used. Spinal cord lysates were first purified using a previously described myelin purification protocol (*34*). Western blots of purified myelin membranes showed significantly decreased expression of mutant human SOD1 in Sox10-Cre; loxSOD1^G37R^ mice when compared to loxSOD1^G37R^ mice, lowering the level to 45% of control when normalized to total protein levels (Fig. 2D and 2E). SOD1 was not detected in the purified myelin membranes of WT controls or Sox10-Cre controls (Fig. 2D), due to the low abundance of wildtype mouse SOD1 in purified myelin membranes. Although less than complete, the 55% reduction in mutant SOD1 is similar to the reduction observed in prior studies of loxSOD1^G37R^ mice (range: 20% to 80% of control) (*4, 5, 21*). Taken together, these data show that Sox10-Cre recombination in Sox10-Cre; loxSOD1^G37R^ mice decreased the abundance of mutant SOD1 in grey matter and white matter of lumbar spinal cord as well as in purified myelin membranes.

**Figure 2.**
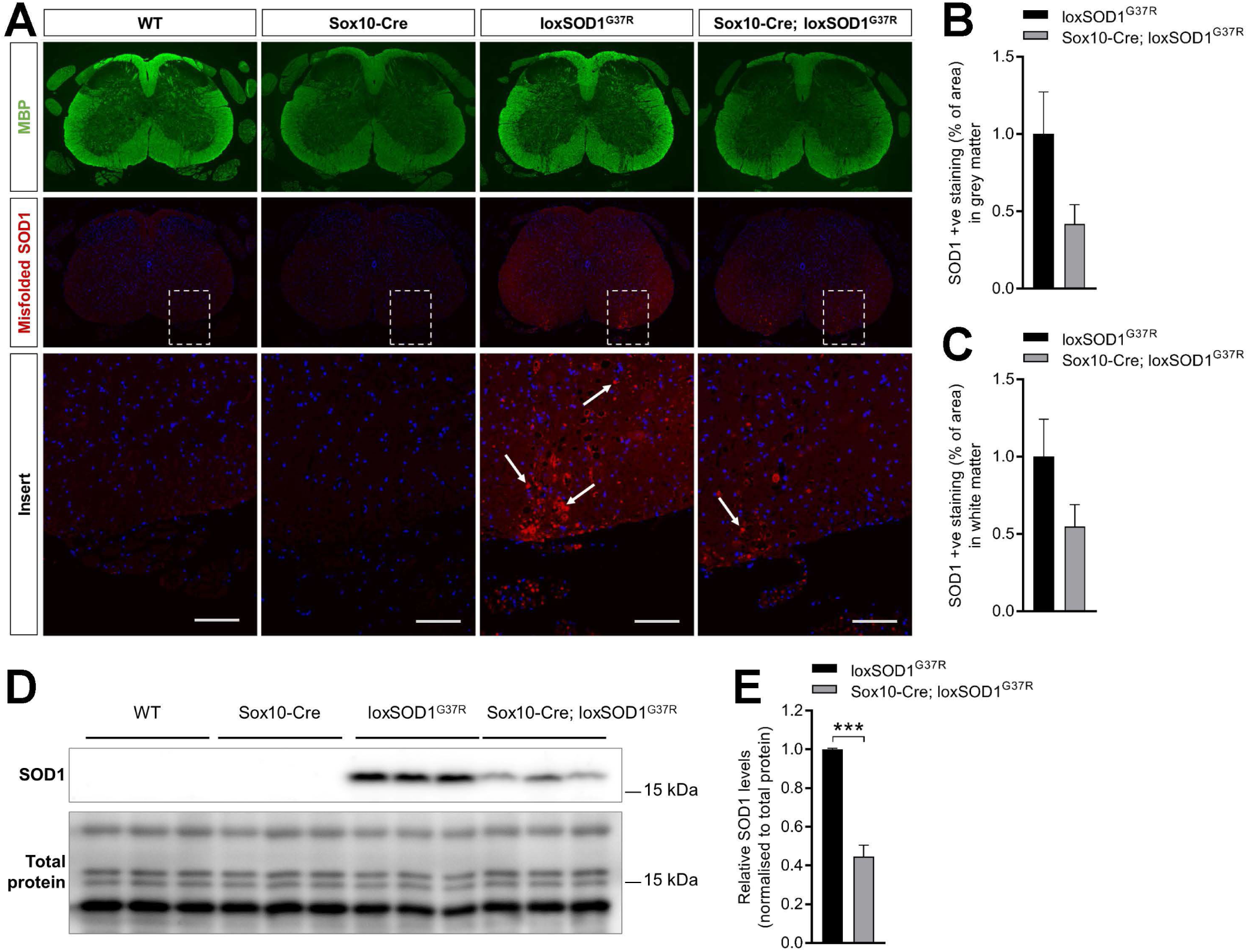
Early removal of mutant *SOD1* expression from oligodendroglial lineage cells decreased mutant *SOD1* expression in spinal cord. A) Immunohistochemistry staining of 44 week old mice showing decreased misfolded SOD1 +ve staining (insert arrows) in lumbar spinal cord of Sox10-Cre; loxSOD1^G37R^ mice when compared to loxSOD1^G37R^ mice. MBP staining is shown in green, misfolded SOD1 staining is shown in red and nuclear DAPI staining is shown in blue. Scale bars represent 100µm. B) Quantification of misfolded SOD1 +ve staining (% of area) in grey matter of immunohistochemistry data shown in A (mean decrease of 58%), loxSOD1^G37R^ (n=5), Sox10-Cre; loxSOD1^G37R^ (n=4), p=0.1150, student’s t test. C) Quantification of misfolded SOD1 +ve staining (% of area) in white matter of immunohistochemistry data shown in A (mean decrease of 45%), loxSOD1^G37R^ (n=5), Sox10-Cre; loxSOD1^G37R^ (n=4), p=0.1767, student’s t test. D) Western blot of purified myelin membranes from spinal cord of 44 week old mice showing decreased expression of mutant human SOD1 in Sox10-Cre; loxSOD1^G37R^ mice when compared to loxSOD1^G37R^ mice (mean decrease of 55%). E) Quantification of mutant human SOD1 levels of western blot data shown in D, loxSOD1^G37R^ (n=3), Sox10-Cre; loxSOD1^G37R^ (n=3), ***p<0.001, student’s t test. Data in B, C and E are expressed as mean values ± SEM.

### Early removal of mutant *SOD1* expression from oligodendroglial lineage cells improves survival and motor performance in ALS mice

To investigate the effect of early removal of mutant *SOD1* expression from oligodendroglial lineage cells on survival, mice were assessed according to established protocols that define disease onset by peak body weight, early disease by a 10% drop in body weight and end-stage by an inability to right itself when placed on the side (*5*). LoxSOD1^G37R^ mice had a median survival of 378 days while Sox10-Cre; loxSOD1^G37R^ mice had a median survival of 448 days (Fig. 3B), indicating that early removal of mutant *SOD1* expression from oligodendrocytes significantly improved survival by 70 days (p<0.0001, Log-rank test). A significant improvement was seen at all stages, including disease onset and early disease (Fig. 3C). Disease onset was delayed by 84.8 days in Sox10-Cre; loxSOD1^G37R^ mice (mean disease onset of 301.6 days) when compared to loxSOD1^G37R^ mice (mean disease onset of 216.8 days). Likewise, the time to early disease was delayed by 116 days in Sox10-Cre; loxSOD1^G37R^ mice (mean early disease of 407.9 days) when compared to loxSOD1^G37R^ mice (mean early disease of 291.9 days). Mice were also assessed for a neurological score, using markers of disease progression established for loxSOD1^G37R^ mice, including hindlimb tremor, hindlimb gait abnormalities (ataxia), hindlimb muscle tone changes, and righting reflex (with a scale of 0 to 2, were zero indicates no abnormalities and 2 indicates end-stage). Consistent with the improved survival, Sox10-Cre; loxSOD1^G37R^ mice showed a significantly improved neurological score when compared to loxSOD1^G37R^ mice (Fig. 3D). No clinical abnormalities were observed in either control groups (WT and Sox10-Cre) throughout the observation period. Lastly, mice were assessed for locomotor function using the rotarod test once weekly from the age of 250 days. Although not statistically significant, there was a noticeable improvement in the rotarod performance of Sox10-Cre; loxSOD1^G37R^ mice when compared to loxSOD1^G37R^ mice (Fig. 3E). WT control mice performed slightly better than Sox10-Cre control mice throughout the observation period, but this difference was not statistically significant. Taken together, these data indicate that removal of mutant *SOD1* expression early during oligodendroglial lineage cell development significantly improved all metrics assessed, including time to disease onset, time to early disease, survival, neurological score, and motor performance.

**Figure 3.**
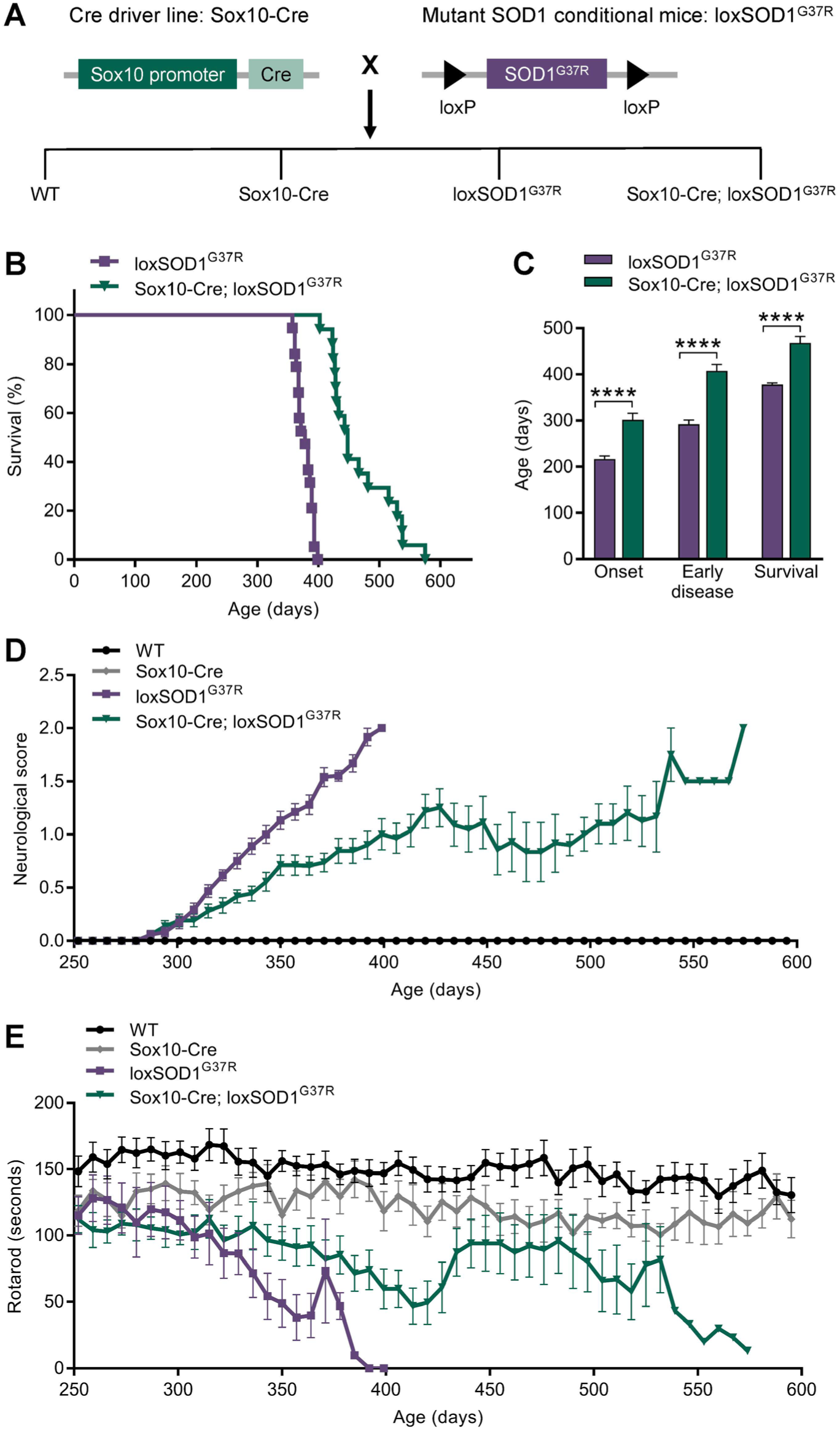
Early removal of mutant *SOD1* expression from oligodendroglial lineage cells improved survival and motor performance in ALS mice. A) LoxSOD1^G37R^ mutant mice were crossed with Sox10-Cre driver mice to remove expression of mutant loxSOD1^G37R^ from Sox10+ oligodendroglial lineage cells. B) Survival to phenotype end-point, median: loxSOD1^G37R^ 378 days (n=19), Sox10-Cre; loxSOD1^G37R^ 448 days (n=17), p<0.0001 Log-rank test. C) Comparison of mean age at disease onset (peak body weight), early disease (10% drop in body weight) and survival, ****p<0.0001, repeated measures two-way ANOVA with Bonferroni post hoc, loxSOD1^G37R^ (n=17), Sox10-Cre; loxSOD1^G37R^ (n=16). D) Assessment for neurological symptoms where a score of 2 indicates end-stage, p<0.0001 when comparing loxSOD1^G37R^ and Sox10-Cre; loxSOD1^G37R^ mice, area under the curve compared using one-way ANOVA with Bonferroni post hoc, WT (n=20), Sox10-Cre (n=17), loxSOD1^G37R^ (n=19), Sox10-Cre; loxSOD1^G37R^ (n=19). E) Performance of mice on the rotarod test for locomotor function, area under the curve compared using one-way ANOVA with Bonferroni post hoc, WT (n=15), Sox10-Cre (n=10), loxSOD1^G37R^ (n=9), Sox10-Cre; loxSOD1^G37R^ (n=10). Data in C, D, and E are expressed as mean values ± SEM.

### Late removal of mutant *SOD1* expression from oligodendroglial lineage cells fails to improve the disease phenotype in ALS mice

Next, we selectively removed mutant *SOD1* expression from mature myelinating oligodendrocytes, using mice expressing Cre recombinase under control of the MOG promoter (hereafter referred to as MOGi-Cre mice; (*35*) that is only active in mature oligodendrocytes (*33*). Crossing hemizygous loxSOD1^G37R^ mice with heterozygous MOGi-Cre mice yielded four genotypes in the expected Mendelian ratio (Fig. 4A), hereafter referred to as wild type (WT), MOGi-Cre, loxSOD1^G37R^, and MOGi-Cre; loxSOD1^G37R^. Unlike the beneficial effects seen with earlier stage deletion of mutant SOD1, no increase in survival was observed when this transgene was deleted at the mature oligodendrocyte stage, with MOGi-Cre; loxSOD1^G37R^ mice exhibiting a median survival of only 370 days (Fig. 4B). Similarly, there was no significant change in time to disease onset or early disease, when comparing loxSOD1^G37R^ mice with MOGi-Cre; loxSOD1^G37R^ mice (Fig. 4C). Time to disease onset differed by only 17.6 days (i.e. not significant) between MOGi-Cre; loxSOD1^G37R^ mice (mean disease onset of 284.5 days) and loxSOD1^G37R^ mice (mean disease onset of 266.9 days), and early disease differed by only 18.1 days (not significant) between MOGi-Cre; loxSOD1^G37R^ mice (mean early disease of 373 days) and loxSOD1^G37R^ mice (mean early disease of 355 days).

**Figure 4.**
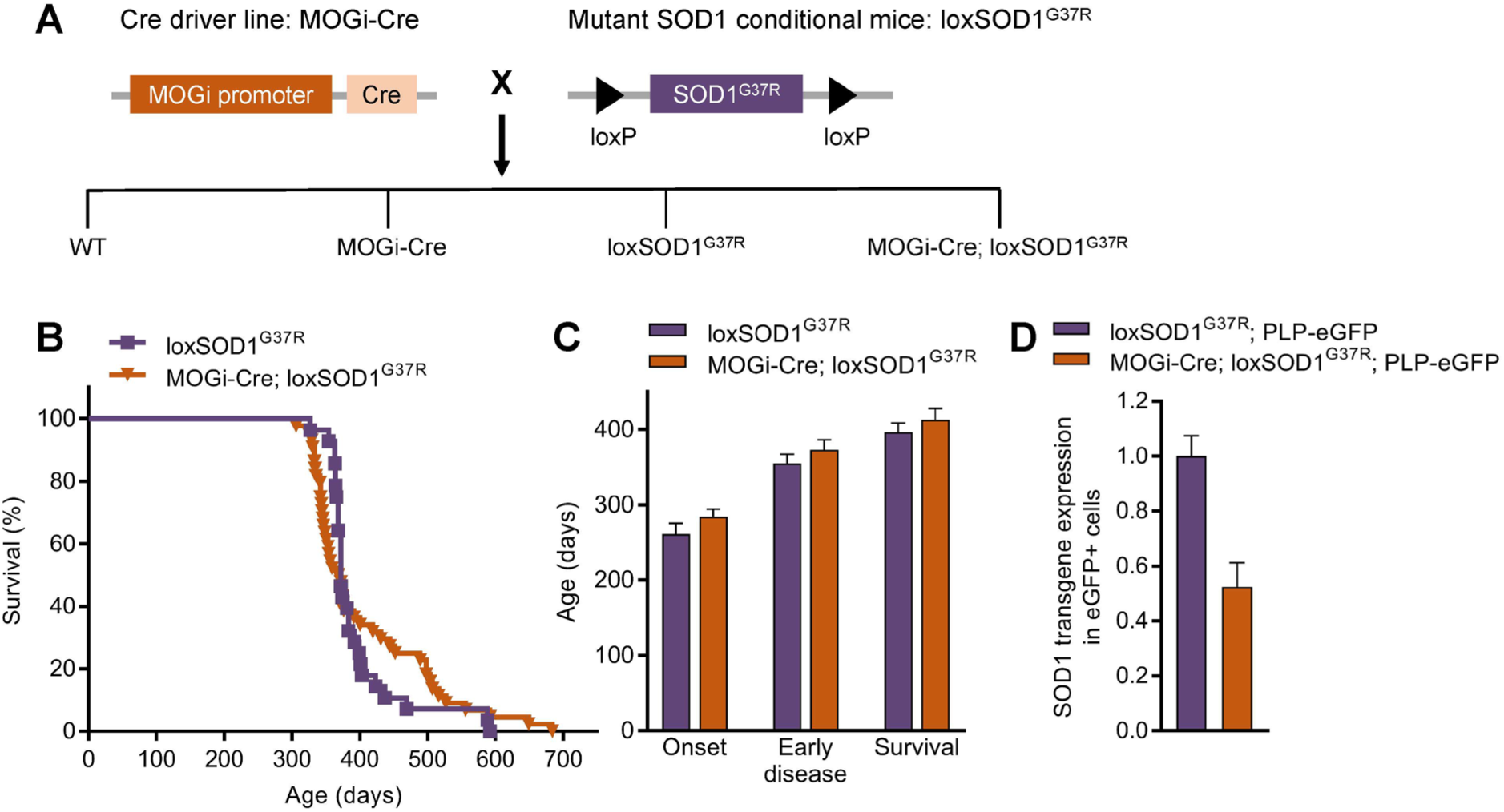
Late removal of mutant *SOD1* expression from oligodendroglial lineage cells did not affect disease onset, early disease, or survival in ALS mice. A) LoxSOD1^G37R^ mutant mice were crossed with MOGi-Cre driver mice to remove expression of mutant loxSOD1^G37R^ from MOGi+ oligodendroglial lineage cells. B) Survival to phenotype end-point, median: loxSOD1^G37R^ 372 days (n=28), MOGi-Cre; loxSOD1^G37R^ 370 days (n=44), p=0.7578, Log-rank test. C) Comparison of mean age at disease onset (peak body weight), early disease (10% drop in body weight) and survival, repeated measures two-way ANOVA with Bonferroni post hoc, loxSOD1^G37R^ (n=26) and MOGi-Cre; loxSOD1^G37R^ (n=41). D) Quantitative PCR of *SOD1* transgene expression in eGFP+ cells FACS sorted from spinal cord of 1.5 month old loxSOD1^G37R^; PLP-eGFP mice (n=2) and MOGi-Cre; loxSOD1^G37R^; PLP-eGFP mice (n=2), mean decrease of 47%. Data in C and D are expressed as mean values ± SEM.

To assess the specificity of mutant SOD1 removal from mature oligodendrocytes, loxSOD1^G37R^ and MOGi-Cre; loxSOD1^G37R^ mice were cross bred with transgenic mice expressing enhanced green fluorescent protein (eGFP) driven by the myelin proteolipid protein (PLP) gene promoter (hereafter referred to as PLP-eGFP reporter mice), which is most active in mature myelinating oligodendrocytes (*33*). To confirm *SOD1* expression was decreased in mature oligodendrocytes in MOGi-Cre expressing mice, all GFP-expressing spinal cord cells from 1.5-month-old loxSOD1^G37R^; PLP-eGFP and MOGi-Cre; loxSOD1^G37R^; PLP-eGFP mice were FACS sorted and subjected to quantitative PCR. *SOD1* transgene expression was significantly decreased (by 47%) in MOGi-Cre; loxSOD1^G37R^; PLP-eGFP mice when compared to loxSOD1^G37R^; PLP-eGFP mice (Fig. 4D). Taken together, these data show that mutant SOD1 was specifically and efficiently excised from mature myelinating oligodendrocytes in MOGi-Cre; loxSOD1^G37R^ mice, and that this did not affect disease onset, early disease, or survival when compared to loxSOD1^G37R^ mice.

### Perturbed myelinic nanochannels worsens the neurological score and survival of SOD1 mutant mice

As we had observed SOD1 aggregates in cytosolic nanochannels of myelin, we aimed to directly test whether a perturbed integrity of myelinic nanochannels can contribute to the pathology and clinical severity of ALS. We crossed SOD1-G93A mutants to mice that lack CNP (CNPnull mice), an oligodendroglial protein that maintains myelinic nanochannel integrity by preventing excessive myelin membrane compaction (*11*). CNPnull mice have a decreased abundance of myelinic nanochannels within the spinal cord as early as 60 days of age (*11*), i.e. long before they develop a clinical phenotype (*36*), and prior to symptom onset in SOD1-G93A transgenic mice. Upon successive cross-breedings we obtained three genotypes, hereafter referred to as CNPnull, SOD1-G93A, and SOD1-G93A; CNPnull (Fig. 5A). Mice were assessed for their neurological score, using established markers of disease progression, including hindlimb tremor, hindlimb gait abnormalities (ataxia), hindlimb muscle tone changes, and righting reflex (*37*). Mice were scored on a scale of 0 to 2, were zero indicates no abnormalities and 2 indicates end-stage. SOD1-G93A; CNPnull mice had a significantly worsened neurological score when compared to SOD1-G93A mice throughout the observation period (Fig. 5B). SOD1-G93A; CNPnull mice also showed a significantly decreased survival (median 175 days) when compared to SOD1-G93A mice (median 245 days), indicating that decreased myelinic nanochannel integrity accelerates disease progression by 70 days (Fig. 5C). Together, these data support the hypothesis that loss of myelinic nanochannels accelerates ALS disease progression by perturbing transport processes from the oligodendroglial to the axonal compartment.

**Figure 5.**
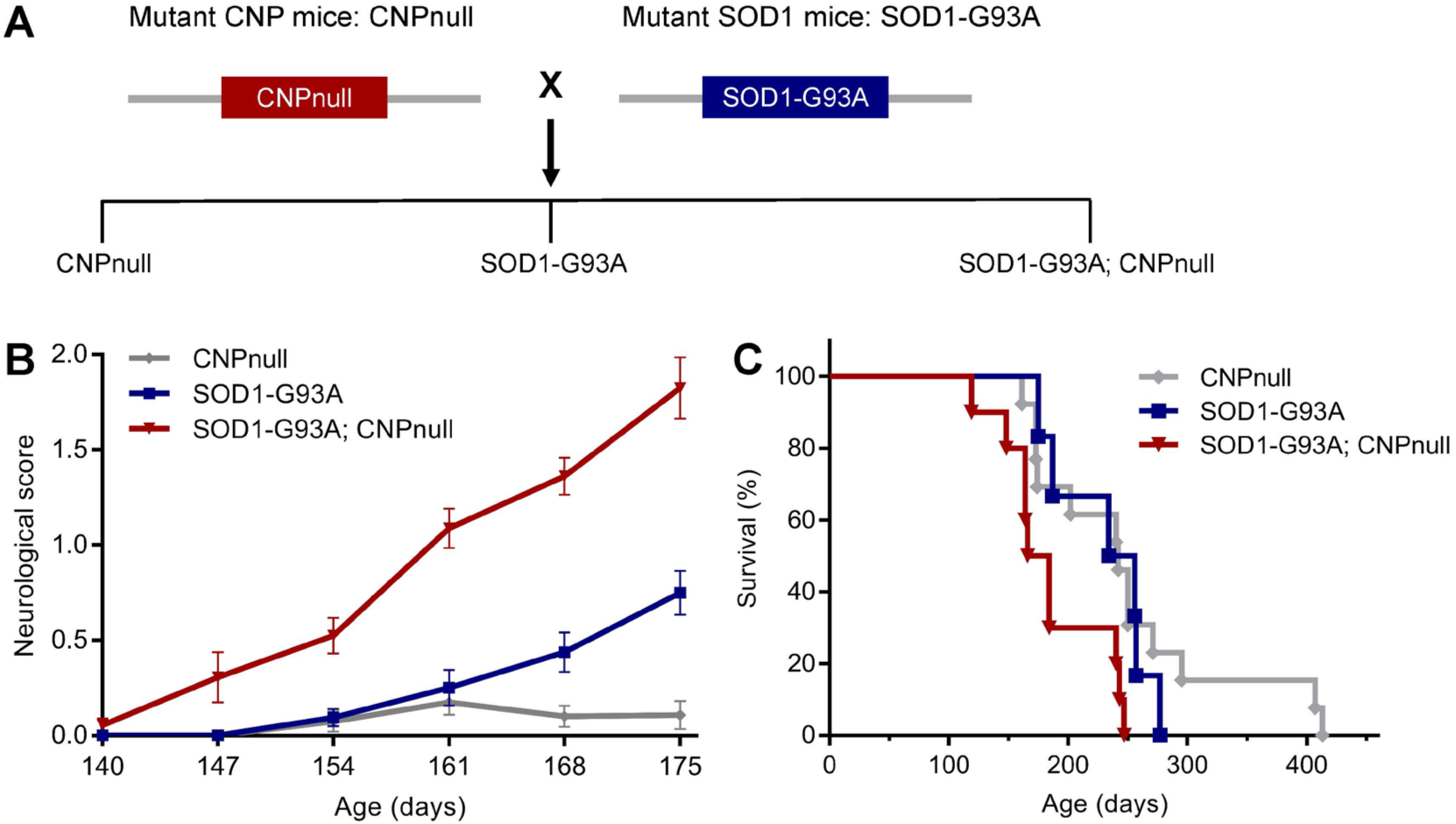
Perturbed myelinic nanochannels worsens the neurological score and survival of SOD1 mutant mice. A) SOD1-G93A mutant mice were crossed with CNPnull mice to decrease the frequency of myelinic nanochannels (successive breedings not shown). B) Assessment for neurological symptoms where a score of 2 indicates end-stage, p<0.0001 when comparing SOD1-G93A and SOD1-G93A; CNPnull mice, area under the curve compared using one-way ANOVA with Bonferroni post hoc, CNPnull (n=10), SOD1-G93A (n=8), SOD1-G93A; CNPnull (n=23). Data are expressed as mean values ± SEM. C) Survival to phenotype end-point, p<0.05 Log-rank test when comparing SOD1-G93A (median 245 days) and SOD1-G93A; CNPnull (median 175 days) mice, CNPnull (n=10), SOD1-G93A (n=6), SOD1-G93A; CNPnull (n=13), CNPnull survival (median 242 days).

## Discussion

The emerging role of oligodendrocytes in ALS pathogenesis has left open the question regarding possible mechanisms by which these cells contribute to disease progression. To experimentally address such mechanisms in the well-known rodent model of SOD1 aggregation and familial ALS, we studied SOD1-G37R mice and selectively removed the mutant *SOD1* gene either early or late during oligodendroglial development. We found that early removal of mutant SOD1-G37R expression delayed disease onset, prolonged survival, and improved motor performance, while late removal did not affect disease onset, early disease, or survival. This led us to hypothesize that accumulation of mutant protein aggregates within myelinic nanochannels could disrupt motor-driven cargo transport and possibly even free diffusion of metabolites from the oligodendroglial soma to the axonal compartment by impairing transport within these nanochannels. This could explain why early removal of oligodendroglial mutant SOD1 is protective, while late removal is not, assuming mutant protein aggregates have already “clogged” myelinic nanochannels. Indeed, we were able to identify mutant SOD1 within these cytosolic channels *in vivo* as well as in cultured oligodendrocytes and their channel-like cellular processes. Furthermore, SOD1-G93A mice with more severely perturbed myelinic nanochannels (SOD1-G93A; CNPnull double mutants) showed a further reduction in survival, indicating that blocking the myelinic nanochannels of oligodendrocytes accelerates ALS as a neuronal disease. Thus, we propose that ALS disease progression is accelerated by reduced transport within oligodendroglial myelin compartments. In support of this model, preventing mutant *SOD1* expression after myelination did not prevent the aggregation of SOD1 that had already accumulated in these spaces.

Considerable evidence indicates that mutant SOD1 contributes to disease pathogenesis of familial ALS through a genetically dominant toxic gain-of-function effect (*38*). This was first confirmed in mice, in which mutant SOD1 was expressed on an otherwise wildtype genetic background, i.e. with two copies of the intact wildtype *Sod1* gene. Cell type-specific effects were later defined by selectively removing a ubiquitously expressed but floxed mutant *SOD1* transgene from specific CNS cell types, in order to better understand the role of that cell type in ALS pathogenesis (*4–6, 21*). Not surprisingly, an improved survival of 64 days was seen when mutant SOD1-G37R was selectively removed from motor neurons (*5*). Importantly, also astrocytes contributed as evidenced by 60 days of improved survival after Cre recombination (*4*), microglial recombination improved survival even by 99 days (*5*). Thus, while ALS is characterized by progressive motor neuron loss, these genetic studies confirmed non-cell autonomous effects of glial cells in ALS pathogenesis. Given the recently discovered role of oligodendrocytes in supplying nutrients to neurons and the reduction of the monocarboxylate transporter MCT1 from oligodendrocytes in ALS mice and human patients (*8*), aggregates of mutant SOD1 in myelinic channels are a plausible cause for the progressively impaired neuronal support. Since motor neurons have high energy demands, even small decreases in the ability of oligodendrocytes to provide nutrients might become problematic over time.

A previous study described SOD1 aggregates in the somata of cultured oligodendrocytes (*24*), which is compatible with our discovery of much smaller SOD1 aggregates in myelinic nanochannels. Our working model is also supported by the finding that removing SOD1-G37R expression from OPCs (*21*), using the Pdgfra-CreER driver line (with two tamoxifen injections at P18 and P30), prolonged survival by 135 days. Interestingly, that rescue was even more efficient than the prolonged survival when the same transgene was removed by constitutive Sox10-Cre excision (70 days). We are unable to adequately explain this quantitative difference, but Sox10-Cre (and not Pdgfra-CreER) is also abundantly expressed in Schwann cells (*39*). Indeed, it was previously reported that selective removal of the same *SOD1-G37R* transgene from Schwann cells decreased (rather than prolonged) the survival of ALS mice by 42 days (*40*). Here, it had been speculated that transgenic overexpression of the enzymatically active SOD1 dismutase in Schwann cells exerts a protective effect on motor axons, possibly by relieving oxidative stress, a rescuing effect that is lost following Cre recombination. It is also possible that mouse husbandries in the US and Europe have developed disease-relevant differences of SOD1-G37R mice, such as the microbiota or the genetic background within inbred substrains (*41*).

Our finding that targeting *SOD1* expression early in oligodendrocyte development improved survival (but late removal did not) is consistent with the observation that the knockdown of SOD1-G37R expression in cultured OPCs (but not in mature oligodendrocytes) reduces the death of co-cultured motor neurons (*24*). Notably, only SOD1-G37R knockdown in OPCs is followed by a later increase in neurotrophic lactate release.

To further investigate the role of perturbed myelinic channel integrity in ALS disease progression, we crossed mutant SOD1-G93A mice with mice lacking expression of CNP, a protein that normally keeps myelinic channels open by preventing excessive MBP-dependent myelin compaction (*11*). SOD1-G93A developed disease at age 5-6 months when CNPnull mice were hardly affected. Double mutants showed earlier ataxia, hindlimb paralysis and end stage disease, indicating that a loss of myelinic nanochannels accelerates ALS as a neuronal disease.

In conclusion, we show that preventing expression of mutant SOD1 early during oligodendrocyte development improved survival, while late removal of this mutant protein did not. A plausible mechanism explaining this finding is that SOD1 protein aggregates form early within myelinic nanochannels, where they impair the ability of oligodendrocytes to provide metabolic and other support to myelinated motor neurons. This mechanism may apply to other diseases in which aggregation prone proteins are expressed by and accumulate in myelinating oligodendrocytes.

### Limitations of this study

Our resulting working model of a detrimental role of aggregation prone proteins in myelinic nanochannels comes from an experimental mouse model that differs from human ALS in many ways, most obviously in the higher expression level of the SOD1 disease gene in mice, but also the slower time course of disease manifestation in humans. Future studies of post-mortem human ALS samples would help reveal whether protein aggregates are also observed within myelinic nanochannels around motor neuron axons.

## Materials and Methods

Full details of the materials and methods are presented in the SI Materials and Methods. Animals used for experiments shown in Figures 1, 2, 3, and 5 were housed at the Max Planck Institute for Multidisciplinary Sciences (City Campus) animal facility and all experiments were conducted in compliance with protocols approved by the local German authorities (Niedersächsisches Landesamt für Verbraucherschutz und Lebensmittelsicherheit). Animals used for experiments shown in Figure 4 were housed at the Johns Hopkins University animal facility and all experiments were conducted in compliance with protocols approved by the Animal Care and Use Committee at the Johns Hopkins University.

## Acknowledgments

Supported by an ERC Advanced Grant (MyeliNANO [KAN]), NIH grants (R01NS033958 [JDR]; R01NS099320 [JDR, BMM]), Muscular Dystrophy Association, and Target ALS. We thank Kathrin Kusch for the MBP antibody. We thank Ramona Jung, Ulli Bode, Caro Boehler, Gudrun Fricke-Bode, Ursula Kutzke, Annette Fahrenholz, Boguslawa Sadowski, Torben Ruhwedel, Nadja Hoffmeister, Conny Casper, Lyudmila Malmaldova, Svetlana Vidensky, and Fang Yang for technical support. We thank members of the Department of Neurogenetics at the Max Planck Institute for Multidisciplinary Sciences and the KAGS subgroup for helpful discussions and input.

## Author Contributions

Designed research: A.I.M., K.A.N., S.G.

Performed research: A.I.M., Y.L., P.D., I.D.T., U.C.G., T.A.B., W.M.

Contributed new reagents, analytic tools and critical advice: D.E.B., B.M.M., J.D.R., D.W.C.

Analyzed data: A.I.M., S.G., W.M., J.M.E., K.A.N.

Wrote the paper: A.I.M., K.A.N.

## Supplementary Information

### SI Material and Methods

#### Mouse models and husbandry

Animals used for experiments shown in Figures 1, 2, 3, and 5 were housed at the Max Planck Institute for Multidisciplinary Sciences (City Campus) animal facility and all experiments were conducted in compliance with protocols approved by the local German authorities (Niedersächsisches Landesamt für Verbraucherschutz und Lebensmittelsicherheit). Animals used for experiments shown in Figure 4 were housed at the Johns Hopkins University animal facility and all experiments were conducted in compliance with protocols approved by the Animal Care and Use Committee at the Johns Hopkins University. SOD1-G93A (*1*), loxSOD1^G37R^ (*2*), Sox10-Cre (*3*), MOGi-Cre (*4*), CNPnull (*5*), and PLP-eGFP (*6*) mice were generated and described previously. All animals were maintained as heterozygotes or hemizygotes, except for CNPnull mice that were maintained as homozygotes. SOD1-G93A, loxSOD1^G37R^ and PLP-eGFP mice were maintained in a mixed background. Sox10-Cre, CNPnull and MOGi-Cre mice were in a C57BL/6 background before being bred with SOD1-G93A or loxSOD1^G37R^ mice. No gender differences in disease development were observed in our studies nor reported by other investigators (*2, 7, 8*). Animals were excluded from the study if they developed severe fight wounds or dermatitis, requiring their immediate euthanasia. All mice were group-housed in individually ventilated cages and kept under a 12:12 light/dark cycle (Max Planck Institute) or 14:10 light/dark cycle (Johns Hopkins University) with access to food ad libitum. For genotyping, genomic DNA was isolated from tail or ear clip biopsies and subjected to routine PCR methods.

#### Mouse behavioral testing and phenotyping

For experiments involving SOD1-G93A and loxSOD1^G37R^ mice, the time of disease onset, early disease and endstage were defined as the time when mouse body weight reached the peak, body weight declined to 10% of the maximum weight and paralyzed mice could not right themselves within 20 seconds when placed on their side, respectively, as described previously (*2*). LoxSOD1^G37R^ mice were assessed for neurological score on a scale from 0–2 as follows: 0, no impairment; 0.5, slight hindlimb weakness; 1, tremor/ataxia; 1.5, hindlimb muscle tone changes; 2, endstage. SOD1-G93A mice were assessed for neurological score on a scale from 0–2 as follows: 0, no impairment; 0.5, tremor/ataxia; 1, hindlimb muscle tone changes; 1.5, moderate paralysis; 2, endstage. For experiments involving loxSOD1^G37R^ mice, motor performance was evaluated by means of the rotarod test. Mice were placed on a horizontal rod that rotated at an accelerating speed (1 rpm every 10 seconds, up to a maximum of 26 rpm) and the time it took for the mouse to fall off was recorded. Mice were tested at 7-day intervals and each mouse was tested three times per trial and the average time was used.

#### Primary oligodendrocyte cultures and immunocytochemistry

Primary oligodendrocyte cultures were prepared as previously described (*9, 10*). Briefly, postnatal day 0-2 (P0-2) cortices from hemizygous mutant SOD1-G93A mice were dissected, dissociated using Papain, and grown as mixed glial cultures in DMEM (Lonza) supplemented with 10% fetal calf serum (FCS), GlutaMAX (Invitrogen), and Penicillin and Streptomycin (Invitrogen) for 12-16 days. OPCs were isolated using mechanical dissociation followed by 20 min differential adhesion on uncoated petri dishes. Remaining microglia were removed by three rounds of immunopanning on dishes coated with 2.3μg/ml BS-lectin-1 in Dulbecco’s phosphate buffered saline (PBS). Purified OPCs were differentiated for 12 days in Sato’s Medium. Differentiated oligodendrocytes were fixed with 4% paraformaldehyde (PFA) for 10 min, permeabilized in 0.1% Triton X-100, washed with PBS, blocked in a solution containing 10% donkey serum and 0.1% Triton X-100 in PBS, and incubated with primary antibodies directed against CNPase (Sigma-Aldrich Cat# C9743, RRID:AB_1840761) and misfolded human SOD1 (MediMabs Cat# MM-0070-P, RRID:AB_2909641) overnight at +4°C. All primary and secondary antibodies were incubated in a blocking solution containing 10% donkey serum and 0.1% Triton X-100 in PBS. Cells were washed three times in PBS before and after secondary antibody incubation. Cells were coverslipped using Fluoromount G (Invitrogen) mounting media and imaged using confocal microscopy.

#### Immunoelectron microscopy

Immunogold labeling of cryosections was prepared as previously described (*11, 12*). Briefly, spinal cords from 5 month old WT and SOD1-G93A mutant mice were dissected and immersion fixed in 4% formaldehyde and 0.2% glutaraldehyde in 0.1 M phosphate buffer containing 0.5% NaCl. After vibratome sectioning (VT1200S, Leica Microsystems, Wetzlar, Germany) pieces of cortical spinal tract and ventral regions with cell bodies of motor neurons were cryo-protected by infiltration with 2.3M sucrose in 0.1M phosphate buffer, placed on aluminum pins and then frozen in liquid nitrogen. Ultrathin cryosections sections were cut using an UC6 ultramicrotome (Leica Microsystems, Wetzlar, Germany) and a 35° cryo-immuno-diamond knife (Diatome, Biel, Switzerland). Primary antibody directed against SOD1 (Enzo Life Sciences Cat# ADI-SOD-100-D, RRID:AB_2039583) was used. Protein A-gold conjugates were used to allow for the detection of SOD1-gold labelling (Cell Microscopy Center, Department of Cell Biology, University Medical Center Utrecht, The Netherlands). Sections were imaged using a transmission electron microscope LEO912 (Carl Zeiss Microscopy, Oberkochen, Germany) and digital micrographs were taken using a 2048x2048 CCD camera (TRS, Moorenweis, Germany).

#### Immunohistochemistry

For experiments involving loxSOD1^G37R^ mice, 44 week old mice were anesthetized with a lethal dose of avertin and transcardially perfused with ice-cold Hank’s buffered salt solution followed by 4% PFA in 0.1 M phosphate buffer, pH 7.4. Spinal cords were dissected and post-fixed overnight in 4% PFA and phosphate buffer. Fixated spinal cords were subjected to dehydration steps (50% ethanol, 80% ethanol, 100% ethanol, 100% isopropanol, 50% isopropanol and 50% xylol, twice 100% xylol) followed by paraffinization on a STP 120 Spin Tissue Processor (Leica Microsystems). Samples were embedded in paraffin blocks on a HistoStar embedding workstation (Epredia). Paraffin-embedded blocks of lumber spinal cord were sectioned transversely in 5-μm slice thickness. Slices were mounted onto slides and dried overnight. Slides were deparaffinized at 60 °C followed by incubation in xylol (100% twice) and a 1:1 mixture of xylol and isopropanol (once) for 10 min each. The slides were rehydrated in a descending ethanol series (100% ethanol, 90% ethanol, 70% ethanol, 50% ethanol, and water) for 5 min each. This was followed by incubation in sodium citrate buffer (0.01 M, pH 6.0) for 5 min at RT and boiling for 10 min. The samples were cooled for 20 min and washed in Tris buffer with 2% milk powder for 5 min. Subsequently, slides were blocked for 20 min at RT in 20% goat serum (BSA/PBS) and incubated at +4°C overnight with primary antibodies directed against MBP (rabbit, custom-made in the Nave Laboratory, 1:5,000) and misfolded human SOD1 (mouse, MediMabs Cat# MM-0070-P, RRID:AB_2909641, 1:500). Slides were then washed 3 times for 5 min each in Tris buffer with 2% milk powder. Slides were incubated for 1 hour at RT in the dark with DAPI (Thermo-Fisher, 1:20000) and the corresponding fluorescent secondary antibodies: anti-mouse Alexa555 (donkey/goat, Thermo-Fisher; 1:1,000) and anti-rabbit Dylight633 (donkey/goat, Thermo-Fisher; 1:1,000). Slides were again washed 3 times for 5 min each in the dark using Tris buffer. Then slides were mounted with Aqua PolyMount mounting medium (PolySciences) and allowed to dry overnight at +4°C in the dark. Mounted slides were imaged using an epifluorescence microscope (Zeiss Axio Imager Z1) and Zen software (Zeiss) using appropriate excitation and emission filters. Quantification of misfolded SOD1 +ve staining was calculated by performing thresholding and the percentage of positive area was determined using Fiji in the respective ROIs (grey matter and white matter).

#### Myelin purification

Purification of the myelin enriched fraction was performed as previously described (*13*). For experiments involving loxSOD1^G37R^ mice, 44 week old mice were killed by cervical dislocation, full spinal cords were dissected out and fresh-frozen on dry ice. Tissues were homogenized in 0.32 M sucrose with protease inhibitor and carefully layered on top of a 0.85 M sucrose solution and then centrifuged for 30 min at 75,000 × g at +4°C. The interphase, containing crudely purified myelin, was carefully collected, resuspended in water, and centrifuged for 15 min at 75,000 × g at 4 °C. The pellet was subjected to osmotic shock by resuspension in water for at least 10 min followed by centrifugation for 15 min at 12,000 × g at +4°C. After a second osmotic shock performed as above, the pellet was resuspended in 0.32 M sucrose solution containing protease inhibitor, carefully added on top of a 0.85 M sucrose solution, and centrifuged for 30 min at 75,000 × g at +4°C. The interphase containing purified myelin was carefully collected, resuspended in water, and centrifuged for 15 min at 75,000 × g. The pellet was resuspended in 1×TBS (supplemented with protease inhibitor, Complete Mini, Roche), and stored at −80°C.

#### Western blotting

To determine the protein concentration in purified myelin samples, the DC protein assay (BioRad Laboratories) was used according to the manufacturer’s guidelines. Samples were then mixed with sample buffer to a final concentration of 2% (w/v) LDS, 0.125% (v/v) Triton X-100, 0.125% (w/v) sodium deoxycholate, 0.501 mM EDTA, 10% glycerol, 247 mM Tris–HCl (pH 8,4), 0.01875% (w/v) Coomassie G250, 0.00625% (w/v) phenol red, 25 mM DTT and 1 μg/μL total protein and heated at +40°C (with shaking at 750 rpm) for 10 min. 15 μg total protein was loaded per lane and separated on 12% Tris-Glycine SDS-PAGE gels in Tris-Glycine SDS running buffer for 1.5 h at 120 V using the Mini protean chamber (Biorad). Proteins were transferred onto a PVDF Membrane (Immobilon-FL PVDF, IPFL00010, Merck Millipore) in transfer buffer (25 mM Tris, 192 mM glycine, 10% methanol) using a XCell II™ Blot Module (Thermofisher) for 1 h at 38 V. To verify equal loading, blots were stained by fast green (5 mg/L fast green, Sigma, in 6.7% acetic acid and 30% methanol) immediately after transfer for 5 min in the dark, washed twice for 30 sec in 6.7% acetic acid and 30% methanol, and imaged using a ChemoStar fluorescent imager (Intas) equipped with a 670 nm/20 nm excitation filter and near-infrared emission collection. Blots were destained in 50% ethanol in TBS-T (150 mM NaCl, 10 mM Tris/HCl [pH 7.4], 0.05% Tween-20) twice for 5 min each and then washed twice in distilled water for 30 sec each. Membranes were blocked in 5% skim milk in TBS-T for 1 h at RT on a rotating shaker. Membranes were then incubated in primary antibody directed against human and mouse SOD1 (Enzo Life Sciences Cat# ADI-SOD-100-D, RRID:AB_2039583, 1:1000) overnight at +4°C on a rotating shaker. After washing with TBS-T, blots were incubated with the secondary antibody anti-rabbit IgG (H+L)-HRPO MinX none (Dianova, Cat# 111-035-003, 1:5,000) for 2 h at RT on a rotating shaker. After washing with TBS-T, blots were detected using WesternBright Sirius HRP substrate (Advansta, Cat# K-12043) and the Odyssey platform (Licor). For quantification of protein abundance, bands were analyzed using Fiji. Protein levels were normalized to fast green total protein.

#### RNA extraction and quantitative PCR

To determine Cre recombination efficiency *in vivo*, spinal cord neural cells from 1.5-month-old loxSOD1^G37R^; PLP-eGFP and MOGi-Cre; loxSOD1^G37R^; PLP-eGFP mice were dissociated using Miltenyi Dissociation Kit (P) following the manufacturer’s instructions. Myelin was removed by using myelin removal beads (Miltenyi). eGFP+ cells were sorted by fluorescence-activated cell sorting (FACS) using MoFlo MLS high-speed cell sorter (Beckman Coulter) at Johns Hopkins School of Public Health FACS core. Dead cells were excluded by propidium iodide staining during sorting. Genomic DNA was extracted from cultured or FACSed cells using QIAmp DNA micro kit (Qiagen) and quantitative PCR for *SOD1* transgene was performed using the primers and cycler parameters on ABI Plusone cycler, as described previously (*2*).

#### Statistical analysis

The group size (number of animals = n) and the statistical tests used are indicated in the respective figure legend. Graphs represent means, and errors are reported as the standard error of the mean (SEM). To account for missing values due to disease progression in neurological score and rotarod data, area under the curve was calculated for each animal and groups were compared using one-way ANOVA with Bonferroni post hoc. All statistical analyses were performed using GraphPad Prism 7. *P* < 0.05 was considered to be statistically significant.

